# Ecological association between seagrass and mangrove ecosystems increases seagrass population longevity in island ecosystem

**DOI:** 10.1101/707745

**Authors:** Amrit Kumar Mishra, Deepak Apte

## Abstract

We report for first time about tropical seagrass meadows association with mangrove ecosystems and its effect on seagrass population dynamics off India in Andaman Sea. Two sites of Neil island, i.e. site 1, associated with mangroves and site 2 without mangroves were selected. Quadrat sampling (n=5) were used to collect sediment and seagrass samples. Reconstruction techniques were used to derive population age structure. *T. hemprichii* population was found mostly with sandy substrate at both sites, with silt consisting very low fraction at site 1. Density, biomass, productivity and morphometric features of *T. hemprichii* were significantly higher at site1. Reproductive density was higher at site 1, whereas reproductive effort to produce fruits were higher at site 2. The rhizome (vertical+ horizontal) production rates were higher at site 1 and the vertical elongation rate was higher at site 2. Plastochrome interval for site1 and 2 were 25.49 and 26.80 days respectively leading to formation of 14.31 and 13.62 leaves per year. *T. hemprichii* population at site 1 had four years of longevity and higher younger plants compared to site 2. The long-term average recruitment and present recruitment rate were higher at site 1 compared to site 2, resulting in steady state growth of the overall population at site 1. Higher number of younger plants suggests fitness of the *T. hemprichii* population at site 1, which increases the ecological significance of mangrove ecosystems have on seagrass population dynamics. This association should be considered for better management and conservation practices of coastal seascapes under global change scenarios.

## Introduction

The dynamic trio of tropical coastal ecosystems; seagrass, mangroves and coral reefs are ecologically productive, biologically diverse and economically valuable and play a significant role in ecosystem functioning and supporting trophic interactions (Kathiresan & Alikunhi, 2011; Guannel et al., 2016). The combined power of these three coastal seascapes increases the resilience of coastal shore lines to natural disturbances (Guannel et al., 2016; Boudouresque et al., 2016) and supply valuable ecosystem services that contribute significantly to the wellbeing of humans (Nordlund et al.,2017). Out of the three, seagrass ecosystems are one of the diverse and widely distributed submerged coastal angiosperms that inhabit the tropic and temperate bioregions of the world except Antarctica (Spalding et al., 2003; Short et al., 2011). These ecosystems supply 24 different ecosystem services such as providing habitat and nurseries for commercially important fish population and endangered sea cows (Cullen-Unsworth et al., 2018), carbon sequestration and storage (Duarte et al., 2013a), shore line protection from storm surges (Boudouresque et al., 2016), prevention of coastal erosion (Christianen et al., 2017; Potouroglou et al., 2017), regulation of nutrient cycles (Constanza et al., 2014) and acting as bioindicator of coastal pollution (Lewis and Devereaux., 2009; Bonanno et al., 2017). These ecosystem services are critical in functioning of seagrass dependent trophic levels, that directly help millions of coastal communities by supplying livelihood and food security (Nordlund et al., 2017). Though coastal communities across the world reap benefits from seagrass ecosystems, human activities including modification and destruction of coastal habitats have led to decline of 30% of seagrass population (7% year^−1^) worldwide (Pendleton et al., 2012; Howard et al., 2014) and much of this 30% of loss has occurred in the bioregion of Asia in the Andaman Sea, Java Sea, South China Sea and Gulf of Thailand (Short et al., 2011; Saenger et al., 2012).

Saying that, research on seagrass associated mangroves or vice versa and their interactive effects on each other for better ecosystem functioning has increased in the last few decades (Nagelkerken et al., 2008; Kathiresan and Alikunhi, 2011). Both ecosystems are physico-chemically linked at macro-level providing a clear evidence of export of organic matter, nutrients, trace metals from these ecosystems to the surrounding coastal environment (Kristensen et al., 2008; Kathiresan and Alikunhi, 2011). Seagrass ecosystems close to mangrove zonation have shown a increase in percentage cover and biomass, as observed for 10 seagrass species (including *Thalassia hemprichii*) associated with mangrove ecosystems were off Thailand coast in Andaman Sea (Poovachiranon and Chansang, 1994). Seasonality influences the mangrove run-off, which in-term affects the seagrass growth, survivorship and distribution (Lirman and Cropper, 2003). Interconnectivity between seagrass and mangrove ecosystems resulting in significantly higher species richness, fish assemblages, density, diversity and population structure have been reported from the Indo-Pacific region (Nagelkerken et al., 2000; Cocheret de la Moriniére et al., 2002; Dorenbosch et al., 2005,2007), whereas lower fish population structure was observed in Pureto-Rico (Aguilar-Perera and Appledorn., 2008) and mixed response was observed off Hainan Island, China (Wang et al., 2009). Such inter-linkage and habitat connectivity between seagrass and mangroves compliment each other in increasing their own productivity and enhance ecosystem functioning of the surrounding marine communities (Dorenbosch et al.,2007; Medina-Gomez et al., 2016) resulting in increase in biodiversity and plant fitness.

India is one of the megadiverse countries, that has wide distribution of 16 seagrass species around its coastal regions (1-18m depth) including the islands of Andaman and Nicobar (ANI) and Lakshadweep (Ragavan et al., 2016). 13 out of 16 seagrass species of India (third highest), covering an area of 29.3 km^2^ are reported from ANI islands (Nobi et al., 2013; Thangaradjou and Bhatt, 2018; Fortes et al., 2018) surrounded by Andaman Sea. Even though ANI has rich mangrove and seagrass biodiversity, research on benefits of seagrass ecosystem due to the presence of mangroves or vice-versa are rarely explored (Kathiresan and Alikunhi, 2011), because sites where both seagrass and mangroves co-exist within similar hydrodynamic conditions are not easily accessible because of the remoteness of the islands. *T. hemprichii* is one of the widely distributed seagrasses of ANI found in the intertidal muddy flats, sandy regions and coral rubbles up-to a depth of 15m (Jagtap et al., 2003; Thangaradjou and Bhatt, 2018) found as individual patches or mixed patches with *Halophila ovalis* or *Halodule pinifolia* (Ragavan et al., 2016). Similar mixed meadows of *T. hemprichii* and *H. ovalis* and its favourable association with mangroves and coral reefs have also been reported from Thailand coast of Andaman Sea (Chansang and Poovachiranon, 1994). Earlier studies on *T. hemprichii* of ANI include their occurrence (Jagtap et al., 1991; Das,1996), species composition, distribution and diversity (Saxena et al., 2010; Ragavan et al., 2016), density, biomass and morphometrics (Savurirajan et al., 2018), heavy metals content (Nobi et al., 2010) and leaf reddening (Ragavan et al., 2014). However, these studies do not provide evidence of the *T. hemprichii* population structure and growth dynamics.

Globally on the background of severe decline of seagrasses leading to extinction risk of 11 seagrass species (Short et al., 2011; Saenger et al., 2012) understanding the population dynamics of seagrass ecosystems is of utmost importance. In India, seagrass beds are declining too, due to anthropogenic disturbances (Thangaradjou et al., 2009; Nobi et al., 2011) and under this scenario, understanding the population dynamics and current growth status of *T. hemprichii* of ANI can help in managing this bioregion. Keeping in mind the increase in tourism and its negative impacts on seagrass population of ANI (Mishra et al., 2019, accepted article), we surveyed the *T. hemprichii* meadows of Neil island to derive the density, reproductive effort, biomass, morphometrics, growth, production and population structure; hence population dynamics of *T. hemprichii* associated with mangrove ecosystem and compared these results with *T. hemprichii* meadows without mangroves. We will test the hypothesis that being associated with mangroves can result in higher plant growth, higher meadow production, and thus better population dynamics for seagrass ecosystems.

## Methods

We surveyed the Shahid Dweep (Neil island) of Andaman and Nicobar Islands (ANI) of India in March 2019. Neil island is situated in south east region of Andaman and Nicobar Islands (Fig.1) has a tidal amplitude of 2.45m, 26.28 to31.67°C of temperature range and salinity between 32 to 35 ppt. Two sites; one with seagrass beds adjacent to mangroves (*Rhizophora apiculata*) and the second without mangroves were selected of Neil island were selected (Fig.1b,c). The two sites were 1000 m apart and were separated by dead coral patches. The habitat configuration of this site is dominated by mangrove *Rhizophora apiculata* in the high tide range, followed by seagrass beds of *Thalassia hemprichii* and surrounded by dead coral patches in the low tide water mark (Fig.1b). The island is surrounded by dead and few live fringing and barrier coral reefs where they provide a suitable habitat for seagrasses (*T. hemprichii*) and mangroves (Nobi et al., 2010) in the lagoon area as observed in the current study site.

**Fig 1.**
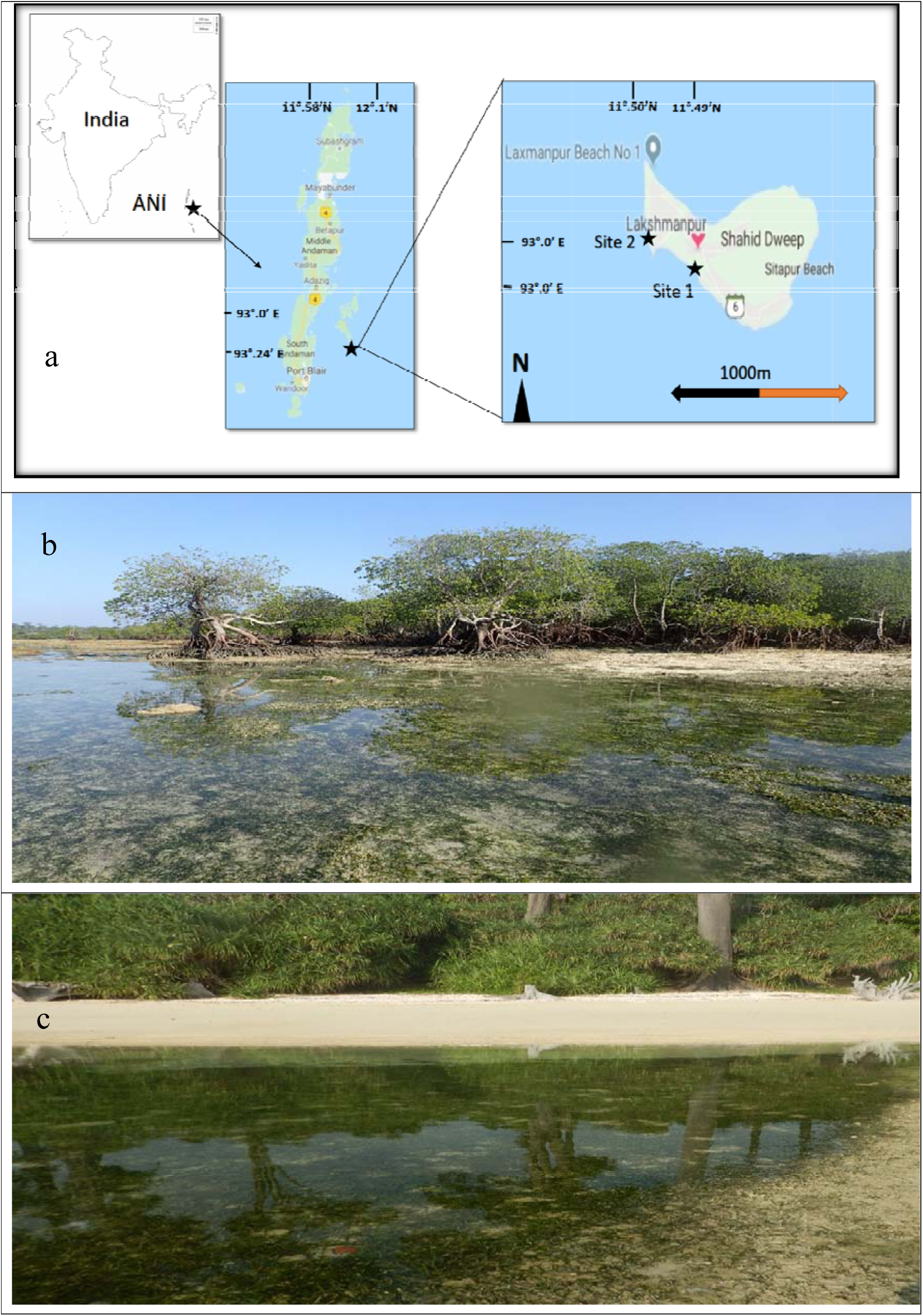
a) Study area showing site 1 and 2 of Neil island of Andaman and Nicobar Islands, India, b) site 1 with mangroves and c) site 2 without mangroves.

Sediment cores (n=5) were collected from each quadrat where seagrass was sampled using 5cm diameter and 10 cm long plastic core. Sediments were collected in plastic bags and were oven dried at 60°C for 72 hours before sieved for grain size fractions.

### Seagrass sampling and analysis

The meadow density and biomass, the rhizome growth and production, the morphometric characteristics of the plants, the population age structure and derived population dynamics (long-term average recruitment, present recruitment and population growth rates) of plants were characterized between the two sites. Reconstruction techniques, an indirect measure of plant growth history and population dynamics changes (Duarte et al., 1994; Fourqurean et al. 2003), were used to evaluate the *T. hemprichii* responses for being associated with and without mangrove ecosystem.

Quadrats (n=5) were collected from two separate sites of *T. hemprichii* within a depth of 0.5m during low tide. We used a quadrat of 20cm^2^ and a hand shovel to dig out seagrass samples up to 10 cm depth. From each quadrat seagrass leaves, rhizomes and roots were collected in plastic bags and brought to the lab for further analysis. The sediment was carefully rinsed off to prevent the modular sets disconnecting from each other and to keep the rhizome mat intact as required for the reconstruction of seagrass dynamics (Duarte et al., 1994). In each sample, the density was estimated by counting the number of both shoots and apicals of physically independent individuals. The presence of reproductive structures, i.e. male of female flowers, in each shoot was recorded to estimate reproductive shoot density and reproductive effort (% of total shoots). Morphometric variables such as horizontal rhizome length, leaf length, width and internode lengths were measured using a Vernier Calliper. The canopy height of *T. hemprichii*, i.e., the leaf length of the longest leaf from the sediment to the leaf tip was measured using a ruler (Mckenzie, 2007). The percentage cover of the seagrass was estimated from the area covered by seagrass from the total quadrat area. The leaves, vertical rhizomes, horizontal rhizomes and roots were separated and dried for 48 h at 60° C for biomass and production estimates.

The age of *T. hemprichii* shoots was estimated by counting the number of leaf scars on the vertical rhizomes plus the number of leaves in each shoot multiplied by the leaf plastochrome interval (PI). To estimate the PI of each study site, i.e., the time needed to produce a new leaf, the sequence of average internodal length of *T. hemprichii* shoots collected with the quadrats plus additional plants collected by hand was plotted. Then a 30% running average was applied to filter short-term seasonal variability and the difference in the number of vertical leaf scars between two consecutive length modes was counted. The modes represent annual growth periods and thus the average number of leaf scars produced between modes was averaged to estimate the leaf PI of the population (Short et al. 2001).

To estimate the vertical and horizontal rhizome elongation rates the length of both the vertical and horizontal rhizomes between consecutive shoots was measured and the number of both vertical and horizontal internodes between consecutive shoots was counted (see Duarte et al., 1994 for details of method). The number of leaves per shoot were measured from intact shoots in each sample (n >150). The horizontal and vertical rhizome production rates were estimated by multiplying the elongation rates (vertical or horizontal) by density (shoots or apicals) and by the specific dry weight of rhizomes (vertical or horizontal).

The long-term average recruitment (R) was estimated from the shoot age structure using the general model: N_x_ = N_0_ *e*^−Rx^, where N_x_ is the number of shoots in age class x, N_0_ is the number of shoots recruited into the population; assuming that mortality and recruitment have had no trend over the lifespan of the oldest shoots in the population, i.e. have remained constant over the lifespan of the oldest shoots, with year to year random variation around some mean value of mortality and recruitment (Fourqurean et al. 2003, Cunha and Duarte, 2005). The recruitment for the current year of sampling (R_0_) was estimated using the method described by Duarte et al., (1994). The population growth rate (*r*) was estimated as: *r* = R_0_ – M, where M is the long-term mortality rate, which equals the long-term recruitment rate (R) under the assumptions of near steady state (Fourqurean et al., 2003). Population was considered growing if *r* is positive (R_0_ > R), shrinking if *r* is negative (R_0_ < R), or with the same trajectory pattern if R_0_ is not significantly different from R (Fourqurean et al., 2003).

The species vertical and horizontal rhizome elongation and the population recruitment rates were obtained considering all replicates in each site. The t-test for the difference between two regression lines was used to compare the vertical rhizome elongation rates as these are equal to the slopes of the linear regression between age and size of rhizomes. Statistical analyses were not performed for the horizontal rhizome elongation rate, because just one value was obtained for each site. The confidence limits of the exponential decay regression model used to estimate the long-term average recruitment rate (R) allowed its statistical comparison to the present recruitment rate (R_0_) as described in Fourqurean et al., (2003). Significant differences of the long-term recruitment rate among sites were tested using one-way ANOVA. Significance levels was considered at *p* < 0.05 (Sokal and Rohlf, 2012).

### Statistics

One -way ANOVA was used to test the significant differences between *T. hemprichii* density, biomass, morphometric features, growth and production estimates between the two sites. All data was pre-checked for normality and homogeneity of variance. Data were log transformed when normality and homogeneity of variance was not achieved for raw data. Tukey’s post hoc analysis was used to test the significant differences (p<0.05). Data are presented as mean and standard error (S.E.).

## Results

Sediment grain size analysis indicated both sites with (site 1) and without mangroves (site 2) were sandy, whereas very small fraction (2.86±0.07%) of silt was observed at site 1 (Table 1).

**Table 1.** Results of grain size analysis of sediments, biomass, morphometrics of *T. hemprichii* beds associated with mangroves (site 1) and without mangroves (site 2) of Neil Island of ANI. Mean ± Standard errors are presented. Different letters indicate significant differences between site 1 and 2. P values from one-way ANOVA are presented; bold letter indicate significant differences. Sand (S), Silt (Si), above ground (AB), below ground (BG), vertical rhizome (VR), horizontal rhizome (HR), not significant (ns), (−) not tested for statistical analysis

| Variables |  | Site1 | Site 2 | p value |
| --- | --- | --- | --- | --- |
| Sediment grain size (%) | S | 97.14±6.13<sup>ns</sup> | 99.98±6.81<sup>ns</sup> | 0.829 |
| | Si | 2.86±0.07 | 0.02±0.0 | - |
| No. of leaves shoot<sup>-1</sup> | | 3.98±0.01<sup>a</sup> | 3.80±0.03<sup>b</sup> | **0.001** |
| No. of leaves year<sup>-1</sup> |  | 14.31 | 13.62 | - |
| Leaf length (cm) | | 10.57 ± 0.21a | 12.28 ± 0.18<sup>b</sup> | **0.003** |
| Leaf width (cm) | | 0.76 ± 0.23<sup>a</sup> | 0.62 ± 0.03<sup>b</sup> | **0.001** |
| Canopy height (cm) | | 19.7±0.15<sup>ns</sup> | 19.3± 0.13<sup>ns</sup> | 0.782 |
| Percentage cover (%) |  | 99 | 98 | - |
| Leaf elongation (cm yr<sup>-1</sup>) | | 0.41±0.01<sup>a</sup> | 0.45±0.01<sup>b</sup> | **0.022** |
| No of vertical rhizomes shoot<sup>-1</sup> | | 1.24± 0.03<sup>ns</sup> | 1.30± 0.04<sup>ns</sup> | 0.934 |
| Rhizome length (cm) | VR | 1.46± 0.10<sup>a</sup> | 1.77±0.11<sup>b</sup> | **0.006** |
| | HR | 8.30±0.40<sup>ns</sup> | 8.65± 0.44<sup>ns</sup> | 0.306 |
| Internode length (cm) | VR | 0.50±0.01<sup>ns</sup> | 0.52±0.01<sup>ns</sup> | 0.694 |
| | HR | 0.34±0.00<sup>ns</sup> | 0.34±0.00<sup>ns</sup> | 0.431 |
| Biomass (g DW m<sup>-2</sup>) | AB | 226.30±4.03<sup>a</sup> | 113.74±1.76<sup>b</sup> | **0.04** |
| | BG | 449.90±9.04<sup>ns</sup> | 423.56±5.23<sup>ns</sup> | 0.812 |
| | AB:BG | 0.57± 0.10 | 0.27±0.02 | - |

Density of *T. hemprichii* at both sites were significant and different, exception was apex density. Both shoot (2741.3±97.3 no m^−2^) and apex density (350.43±66.5 no m^−2^) of site 1 were 2.1-fold and 1.5-fold higher than site 2 (Fig.2a). *T. hemprichii* reproductive density was significant and 1.2-fold higher than site 2, while the reproductive effort at site 1 (2.7%) was 1.7-fold lower than site 2 (Fig.2b).

**Fig 2.**
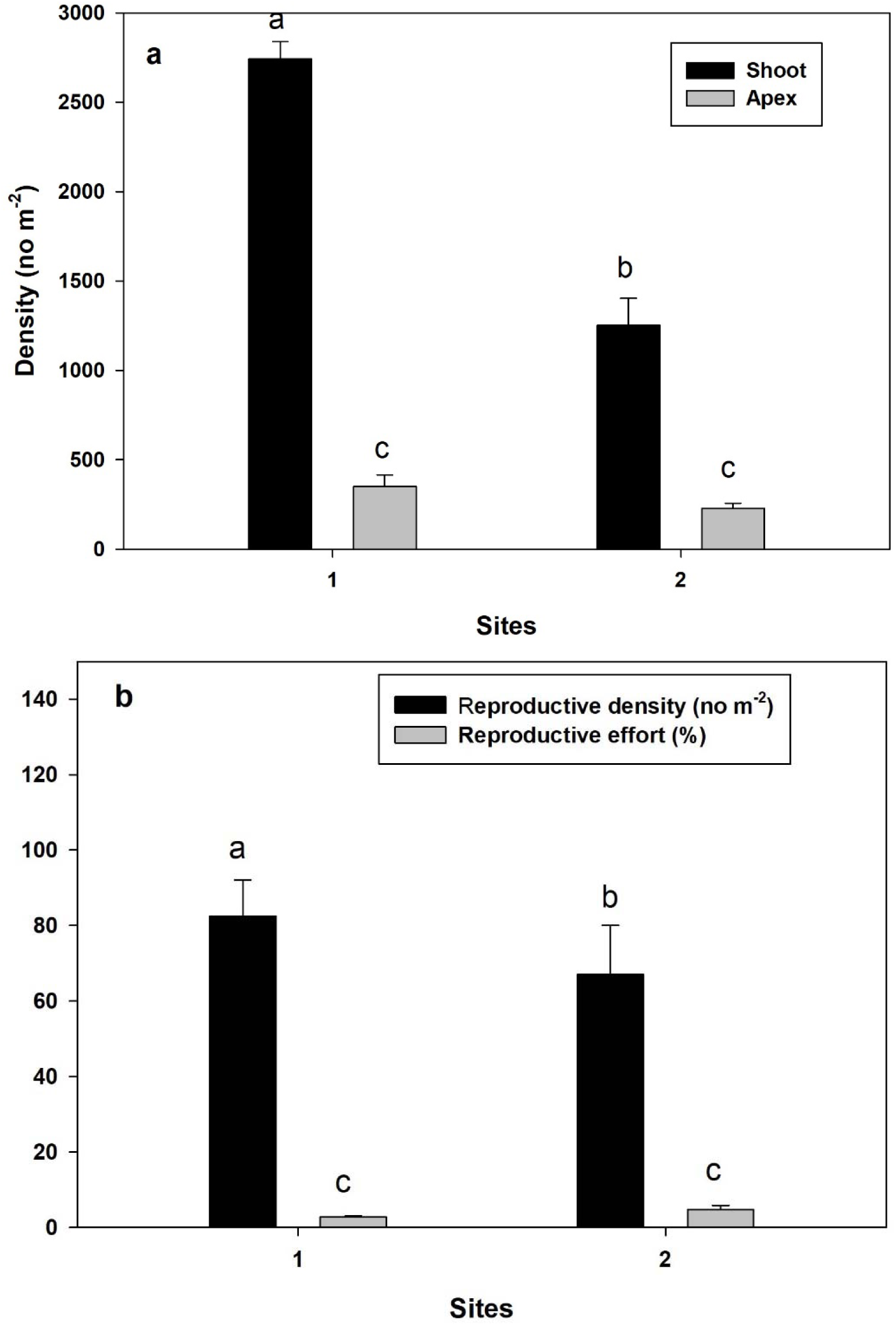
a) Density (shoot and apex) and b) reproductive density and effort of *T. hemprichii* of Neil Island of ANI associated with mangrove (Site 1) and without mangroves (Site 2). Error bars represent standard error. Small letters indicate significant differences between the two sites.

Morphometric features of *T. hemprichii* between the two sites were significant and different for various measurements of leaves and rhizome length, whereas no significant differences were observed for canopy height, number of vertical rhizomes per shoot and internode length (Table 1). The number of leaves per shoot ((3.98±0.01 no shoot^−1^) was higher at site 1 than site 2. Interestingly the average leaf length of *T. hemprichii* at site 1 (10.57 ± 0.21 cm) was 1.1-fold lower than site 2, whereas the leaf width at site 1 was 1.2-fold higher than site 2 (0.62±0.03 cm). However, the canopy height and percentage cover of *T. hemprichii* between the two sites were slightly higher at site 1 compared to site 2. The leaf elongation rate of *T. hemprichii* was 1-fold higher at site 2 compared to site 1 (0.41± 0.01 cm y^−1^). The number of vertical rhizomes per shoot, the rhizome length (both vertical and horizontal) were 1-fold, 1.2-fold and 1-fold higher at site 2 compared to site 1 respectively, whereas the internode lengths of both vertical and horizontal rhizomes were similar between both sites (Table 1).

Biomass of *T. hemprichii* between the two sites were significant and different except for below ground biomass (Table 1). The above ground (AB) and below ground (BG) biomass at site 1 was 2-fold and 1-fold higher than site 2 (AB-113.74± 1.76 g DW m^−2^; BG-423.56± 5.23 g DW m^−2^) respectively. Higher above ground biomass of site 1 contributed to 2-fold higher AB:BG ratio at site 1 than site 2 (Table 1).

The vertical rhizome elongation rates were significantly different between the two sites (Fig.3a). The vertical rhizome elongation rate of site 1 was 1.4-fold lower than site 2 (2.22± 0.11 cm year^−1^), whereas the horizontal rhizome elongation rate of site 1 was 1-fold higher than site 2 (9.58 cm year-1) (Fig. 3a). The rhizome (vertical and horizontal) production rates of *T. hemprichii* between the two sites were significant and different (Fig.3b). Site 1 was recorded with higher vertical and horizontal rhizome production. The vertical rhizome production of site 1 was 1.3-fold higher than site 2 (112.65±19.5 g DW m^−2^ year^−1^), whereas the horizontal production rate of site 1 was 1-fold higher than site 2 (98.94±13 g DW m^−2^ year^−1^) (Fig.3b).

**Fig 3.**
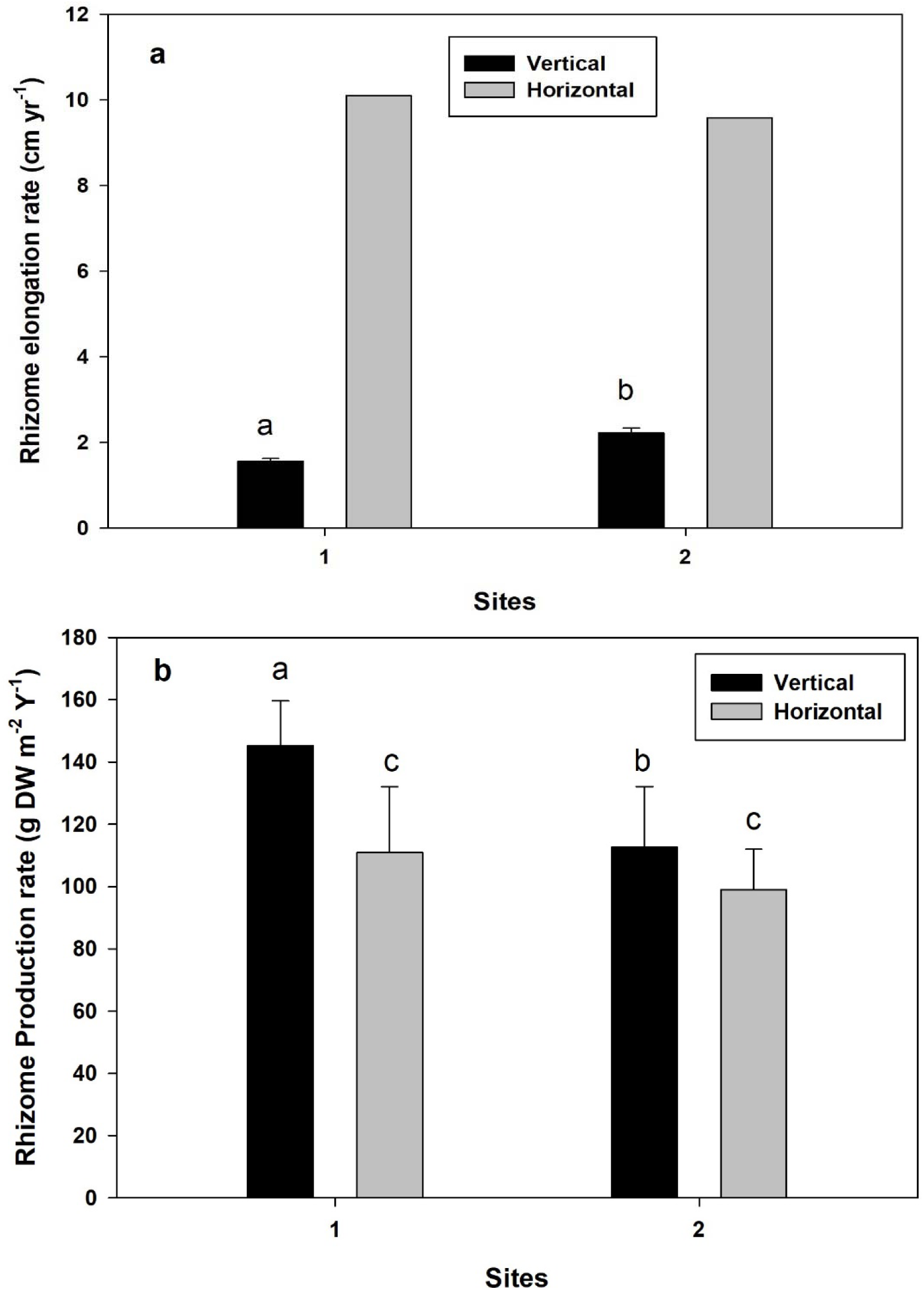
Rates of rhizome (vertical and horizontal) elongation (a) and production (b) of *T.hemprichii* of Neil Island of ANI associated with mangrove (Site 1) and without mangroves (Site 2). Error bars represent standard error. Small letters indicate significant differences between the two sites derived from t-test for vertical elongation rates (p<0.001).

The plastochrome interval of *T. hemprichii*, i.e. the number of days to produce one leaf was different for both sites, with 25.49 days for site1 and 26.78 days for site 2 (see supplementary data), that resulted in production of 14.31 and 13.62 leaves per year (Table 1). Interestingly, the average shoot age of *T. hemprichii* at site 1 (0.85±0.03 year) was lower than site 2 (0.88±0.04 year), whereas the longevity of shoots at site 1(3.30 years) was 1.1-fold higher than site 2 (Table 2). No significant differences were observed for long-term average recruitment (R) between both sites (Table 2). However, R of site 1 was 1.3-fold higher than site 2 (0.85±0.29 year^−1^). The present recruitment (Ro) rate of *T. hemprichii* population sampled for the current year at site 1 was 1.2-fold higher than site 2 (0.97 year^−1^). However, the difference between R and Ro for site 1 was 1.5-fold lower compared to the difference between R and Ro for site 2 (0.12), resulting in higher population growth of *T. hemprichii* at site 2 (0.12 year^−1^) than site 1(0.08 year^−1^) (Table 2). Contrastingly, the age frequency distribution of *T. hemprichii* showed higher younger plants (n=167, of age class one) at site 1 compared to site 2 (n=100) (Fig.4)

**Table 2.** Age structure and population dynamics of *T. hemprichii* shoots from Site 1 (with mangroves) and Site 2 (without mangroves) of Neil island of ANI. Mean ± Standard errors are presented for the shoot age. The exponential coefficient ± standard errors of the exponential decay regression are presented for the long-term average recruitment rate (R). Different letters indicate significant difference between site 1and site 2 only for R, ns = not significant.

|  | Site1 | Site 2 |
| --- | --- | --- |
| Shoot longevity (years) | 3.30 | 2.84 |
| Shoot age (years) | 0.85 ± 0.03<sup>ns</sup> | 0.88 ± 0.04<sup>ns</sup> |
| Long term avg. recruitment rate (R, year<sup>-1</sup>) | 1.11 ± 0.16<sup>ns</sup> | 0.85 ± 0.29<sup>ns</sup> |
| Present recruitment rate (R<sub>o</sub>, year<sup>-1</sup>) | 1.19 | 0.97 |
| Population growth rate (r, year<sup>-1</sup>) | 0.08 | 0.12 |

## Discussion

The ecological-interlinkage between seagrass ecosystems with mangroves are beneficial to each other and play an important role in the functioning of coastal seascapes (Dorenbosch et al.,2007; Medina-Gomez et al., 2016). We observed seagrass (*T. hemprichii*) association with mangroves (*R. apiculata*) resulted in increased density, biomass, morphometrics, recruitment rates and longevity of plants, i.e., increased population dynamics compared to the site without mangroves of Neil island of ANI.

The sediment grain size fraction at both sites were similar with higher percentage of sand content, whereas site 1 associated with mangroves was observed with higher silt content (Table 1), which indicates about influx of fine grain size fractions from mangrove ecosystem (Kathiresan and Alikunhi, 2011) that were detained in the seagrass ecosystem. Saying that, *T. hemprichii* is mostly found within coral rubbles with sandy sediments due to weathering of dead coral rubbles (Tussenbroek et al., 2006; Ragavan et al., 2016). The higher fine sediment content at site 1 can also be a result of sediment filtering and deposition capacity of seagrass, that reduces seawater turbidity above the seagrass canopy (Boer, 2007; Short et al., 2007; Potouroglou et al., 2017) and help in primary productivity and plant growth. Percentage of sand (97-99%) and silt (2.8%) content in the sediment grain size fraction at both sites (Table1) were within similar range (sand-82-98.1%; silt-0.98-15.9%) obtained previously for Neil island (Nobi et al., 2010) and for *T. hemprichii* meadows (sand-86%; silt-9.74%) of ANI (Savurirajan et al., 2018). The higher silt content reported by Nobi et al., (2010) and Savurirajan et al., (2018) are for various mangrove associated or estuarine sites of ANI, which coincides with our results of higher silt content at site 1 with mangroves. The sand and silt content in our studies were similar to *T. hemprichii* meadows around Andaman Sea (Rattanachot et al., 2018)

Total density (shoot + apex) was higher at site 1 compared to site 2 (Fig. 2a), which indicates better growth rates of *T. hemprichii* when associated with mangroves. Total density in our results were 4.6-fold (site1) and 2-fold (site2) higher than previously obtained results for ANI (Savurirajan et al., 2018). This contrasting differences in density can be due to the selection of various sites of ANI by the authors, that are anthropogenically disturbed or modified for various human activities (Sahu et al.,2015) such as aquaculture expansion, leading to loss of *T. hemprichii* biodiversity, whereas both sites of Neil island are more isolated from direct impacts related to tourism and recreational activities. Saying that, *T. hemprichii* total density measurements from Philippines (1358-1853 no m^−2^) around Andaman Sea (Rollon et al., 2001) were lower than site 1 (2741.63 no m^−2^) and higher than site 2 (1252.30 no m^−2^). The lower density of *T. hemprichii* at site 2 can be a result of increased sand wave dynamics resulting in shoot mortality or damage, which has been observed for seagrass population under sediment burial and erosion conditions (Cabaço et al., 2008; Saunders et al., 2017), which has been observed for *T. hemprichii* population of Philippines in Andaman Sea (Duarte et al., 1997). The higher density at site 1 is due to low mortality/physical damage of *T. hemprichii* plants, as they are sheltered from direct impact of burial and erosion by presence of mangrove ecosystems (Saenger et al., 2012; Guannel et al., 2016).

The reproductive density at site 1 was higher than site 2 (Fig.2b), whereas the reproductive effort was higher at site 2, which indicates about the increased reproductive effort by *T. hemprichii* population to physical disturbances caused by the wave dynamics and sand erosion at this site. Higher sedimentation from wave breaking results in breakage of *T. hemprichii* fruits/flowers, leading to higher reproductive effort (McMahon et al., 2017), as the current sampling season (March 2019) is favourable for production of fruits in *T. hemprichii* (Tongkok et al., 2017). Secondly, intertidal exposure to high temperature and light plays a significant role in determining the phenology of seagrasses (Smith and Walker, 2002; McMahon et al., 2017), which in our case is the exposure of *T. hemprichii* plants to low tides twice in a day with high temperatures above 30°C, resulting in reduction of *T. hemprichii* fruit production. Similar results of high reproductive effort and low fruit production were observed for *T. hemprichii* population of southern Thailand (Tongkok et al., 2017) and Indonesia (McMahon et al., 2017) in Andaman Sea. We report here for the first time about reproductive density of *T. hemprichii* for ANI and India.

The number of leaves per shoot and the number of leaves produced per year (Table 1) derived from plastochrome interval for site 1 (25.49 days per leaf) was higher compared to site 2 without mangroves, which suggests the *T. hemprichii* meadows being associated with mangroves have the requisite nutrients and favourable conditions to produce a single leaf within 25.49 days, whereas the site without mangroves is deprived of nutrients as the waters of Andaman Sea are oligotrophic (Mishra and Mohanraju, 2018; Mishra and Kumar, 2019 accepted article) and are under immense pressure from breaking waves. The number of leaves per shoot at site 2 (3.80±0.03) in our studies are similar (3.80) and higher at site 1 (3.98±0.01) to previously observed results of ANI (Savurirajan et al., 2018).

Higher leaf length (12.28 ± 0.18 cm) of site 2 was observed with positive related to increased leaf elongation rate (0.45±0.01 cm y^−1^) compared to site 1, which clearly indicates about the immense pressure on *T. hemprichii* to avoid being buried by sand at site 2. Supporting this trend of increased vertical growth of *T. hemprichii* were also the number of vertical rhizomes per shoot (1.30± 0.04), the vertical rhizome length (1.77±0.11cm), vertical internodal length (0.52±0.01cm) and vertical elongation rate (2.22±0.11 cm y^−1^) which were higher at site 2 compared to site 1. Increase in vertical internodal growth is an essential mitigation feature of *T. hemprichii* to avoid sediment burial, which has been observed for *T. hemprichii* meadows globally (Cabaço et al., 2008).

The mean horizontal rhizome length at site 2 was also higher compared to site 1 (Table 1), which indicates about the need of spatial migration of the meadow towards a more suitable habitat away from the breaking waves, but this meadow migration was not supported by higher horizontal rhizome growth rate (Fig.3a). This contrasting morphometric feature and rhizome growth rate can be a result of dead coral patches that surround the *T. hemprichii* meadows at site 2 (as observed during the field sampling) and hinder the meadow migration. Contrasting to site 2, site 1 with mangroves were observed with lower horizontal rhizome length (Table 1) but increased horizontal elongation rate (Fig.3a), that can help the *T. hemprichii* meadows to migrate spatially and extent the range of *T. hemprichii* habitat. *T. hemprichii* morphometric features such as leaf length, width and internode (vertical and horizontal) length values observed in our studies (Table 1) were significantly higher than previously obtained values for *T. hemprichii* from ANI (Savurirajan et al., 2018) except for horizontal internode length which were lower, whereas the leaf width (0.76-0.81mm) observed at Philippines were higher than site2 and similar to site 1 (Rollan et al., 2010).

The canopy and percentage cover of *T. hemprichii* at site 1 was higher that site 2 (Table 1), though the average leaf length of *T. hemprichii* at site 1 (10.57 ± 0.21 cm) was lower than site 2 (12.28 ± 0.18 cm), the canopy height was higher at site 1, which indicates about the physical damage caused to the leaf canopy resulting in breakage of leaf tips due to sand wave breakage and presence of these meadows in shallow waters makes it more difficult for the plants.

Biomass followed the similar pattern of density with higher above ground and below ground biomass at site 1 compared to site 2. The higher above ground biomass is a result of higher number of leaves per shoot and year and wider leaves compared to *T. hemprichii* population at site 2. Saying that, the *T. hemprichii* population at site 1 was under an ambient and sheltered environment from wave breaking and sand burial due to presence of mangroves, whereas the site 1 was under immense pressure from these physical disturbances. Presence of soft sediments adjacent to mangrove and higher horizontal growth rate helped the *T. hemprichii* population at site 1 in having a greater spatial migration and increasing below ground biomass, whereas at site 2, the lower below ground biomass is a result of lower horizontal elongation rate and unfavourable dead coral barriers to restrict the spatial growth of the meadow. However, the *T. hemprichii* meadows at site 2 grew deeper that resulted in higher root biomass (see supplementary data) compared to horizontal rhizome biomass. Higher below ground biomass and lower above ground biomass of *T. hemprichii* in our studies followed the similar pattern observed for *T. hemprichii* meadows around ANI (Savurirajan et al., 2018) and off Thailand coast (Poovachiranon and Chansang, 1994) in Andaman Sea. The above ground biomass of *T. hemprichii* at site 1(7-fold) and site 2 (3-fold) higher and the below ground biomass are 6-fold higher than previously observed results for ANI (Savurirajan et al., 2018). The total rhizome (vertical+ horizontal) production rate was higher at site 1compared to site 2 (Fig.3b) which indicates that *T. hemprichii* meadows associated with mangroves can increase their vertical and horizontal rhizome production rate due to higher growth and less disturbances.

The plastochrome interval (Site 1 (25.49 days/leaf), site 2 (26.8 days/leaf)) in our studies for *T. hemprichii* were higher, than observed for *T. hemprichii* population of Philippines coast (9.19 days/leaf) (Rollon et al., 2010). However, when compared to global plastochrome interval of *T. hemprichii* (10.9-21.9 days/leaf) data of Short and Duarte, (2001), the *T. hemprichii* population of ANI required more days to produce a single leaf compared to global data, which can be the impact of low nutrient content in the oligotrophic waters of Andaman Sea (Mishra and Kumar, 2019, accepted article) for *T. hemprichii* population. Secondly our sampling period was in summer season, when more energy is diverted towards *T. hemprichii* reproductive cycle than plant productivity for *T. hemprichii* population of Andaman Sea (Tongkok et al., 2018).

The shoot longevity of *T. hemprichii* population was higher at site 1 compared to site 2, whereas the average shoot age of plants was higher at site 2 (Table 2), which indicates about the continuous growth of *T. hemprichii* shoots to overcome burial, whereas being sheltered by mangroves at site 1, *T. hemprichii* population gets to survive longer for four years while at site 2 only for three years. From the age frequency data, we observed that plant population associated with mangroves has higher number of younger plants (<1 year, 167 nos.) compared to the site 2 (<1 year, 100 nos.) without mangroves (Fig. 4). The reduction in number of younger plants at site 2 is a direct impact of sediment burial due to sand wave breaking in shallow waters (Duarte et al., 1997; Cabaço et al., 2008).

**Fig 4.**
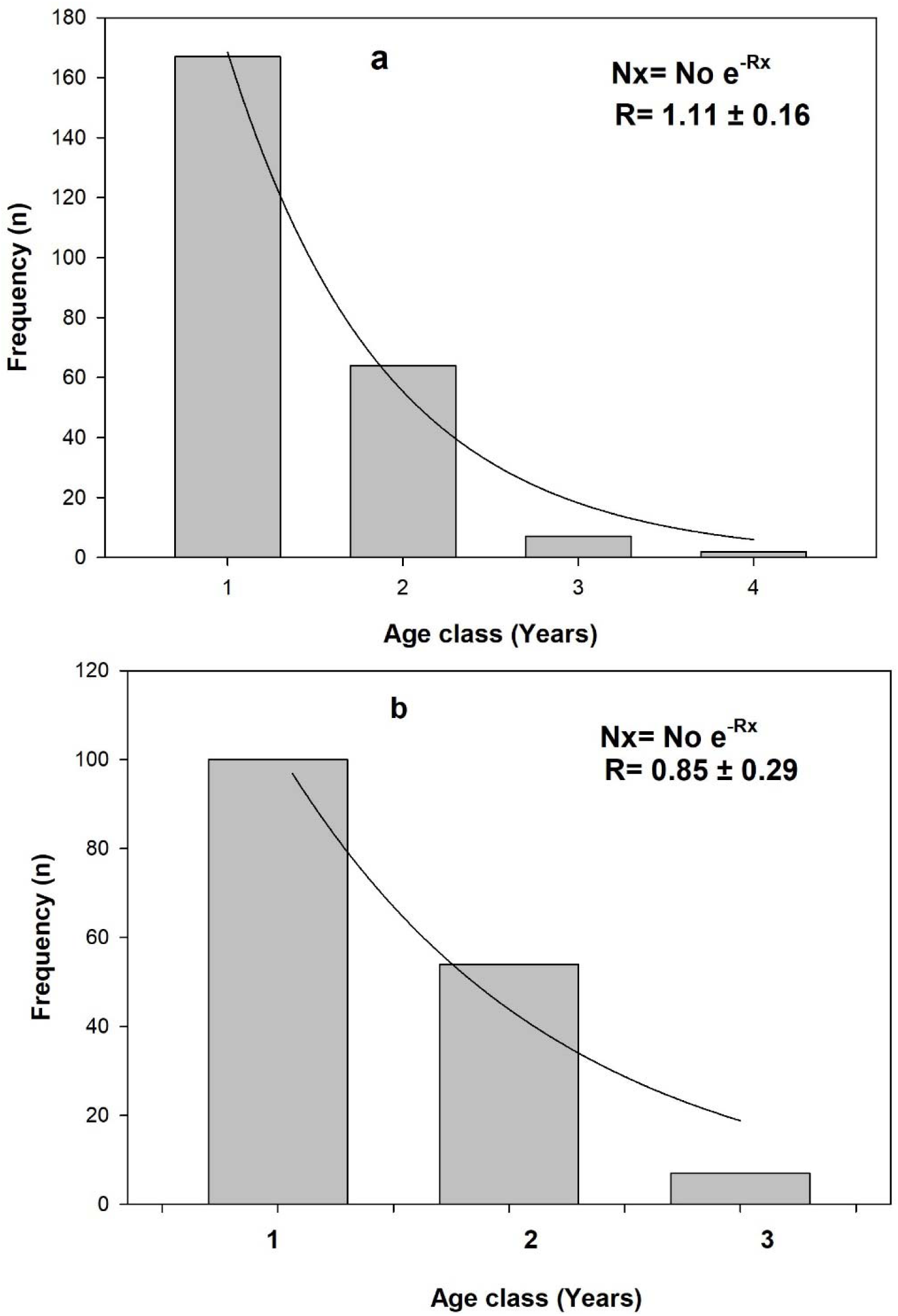
Age frequency distribution of *T. hemprichii* population of Neil Island of ANI associated with mangrove (a, Site 1) and without mangroves (b, Site 2). The long-term average recruitment rate (R) was estimated from the exponential decay regression line fitted to age frequency distribution.

The present recruitment (Ro) was higher than the long-term average recruitment (R) for *T. hemprichii* population at both sites, but R was higher at site 1 compared to site 2. Higher Ro indicates the population is growing in these islands, but the rate of growth (r) is higher for the population under stress at site 2 compared to steady state growth at site 1.Higher Ro and r of site 2 are result of beach dynamics, where *T. hemprichii* meadows are subjected to sand wave breaking and seagrasses being efficient ecosystem engineers are able to change their own environment by sediment filtration or trapping to avoid being buried (Reitkerk et al., 2004) simultaneously stimulating growth of their leaves and rhizomes (Cardoso et al., 2004; McMahon et al., 2017). This higher growth rate can be correlated with higher reproductive effort (4.7%) at site 2, as found in our result compared to site 1 (Fig.2). The population associated with mangroves is more sheltered from increasing wave dynamics by the dead coral patches from the seaward side and from beach dynamics by the presence of mangrove ecosystem, resulting in a steady state growth rate.

## Conclusion

We report for the first time about the population dynamics of *T. hemprichii* population of ANI of India. Seagrass population survives longer when associated with mangroves along with increased density, biomass and morphometrics, suggesting plant fitness, growth and productivity, whereas plants without mangroves has increased leaf length, vertical rhizome elongation rates to survive the burial by sand wave breaking. Reproductive density of plants associated with mangroves were higher compared to sites without mangroves. Growing and healthier *T. hemprichii* population of Neil Islands can provide suitable feeding grounds to green sea turtle populations (Christianen et al., 2017) that navigate these islands along with providing suitable nurseries for marine fisheries (Unsworth et al., 2008). Though our results provide initial data about *T. hemprichii* population dynamics, more studies are required to generate a clear picture of *T. hemprichii* population status along the Andaman and Nicobar group of Islands and other sites of mainland India. This ecosystem interlinkage between seagrass and mangroves should be further investigated, so that the resilience of these coastal seascapes can increase under anthropogenic disturbances and climate change scenarios to have healthier ecosystems that can provide better ecosystem services (Guannel et al., 2016).

## Acknowledgements

We are grateful to the PCCF, wildlife to provide the required official help during the sampling programme.

## References

Aguilar-Perera, A. & R.S. Appeldoorn. 2008. Spatial distribution of marine fishes along a cross-shelf gradient containing a continuum of mangrove–seagrass–coral reefs of southwestern Puerto Rico. Estuarine, Coastal and Shelf Science 76: 378–394

Boer, W.F., 2007. Seagrass-sediment interactions, positive feedbacks and critical thresholds for occurrence: a review. Hydrobiologia. 591, 5–24

Bonanno, G., Borg, J.A., Di Martino, V., 2017. Levels of heavy metals in wetland and marine vascular plants and their biomonitoring potential: A comparative assessment. Science of the Total Environment. 576, 796–806.

Boudouresque, C.F., Pergent, G., Pergent-Martini, C., Ruitton, C., et al., 2016. The necro mass of the *Posidonia oceanica* seagrass meadow: fate, role, ecosystem services and vulnerability. Hydrobiologia. 781, 25–42.

Cabaço, S., Santos, R., Duarte, C.M., 2008. The impact of sediment burial and erosion on seagrasses: A review. Estuarine, Coastal and Shelf Science. 79, 354–366

Cardoso, P. G., M. A. Pardal, A. I. Lillebo, S. M. Ferreira, D. Raffaelli & J. C. Marques, 2004. Dynamic changes in seagrass assemblages under eutrophication and implications for recovery. Journal of Experimental Marine Biology and Ecology 302: 233–248

Chansang, H and Poovachiranon, S., 1994. The distribution and species composition of seagrass beds along the Andaman Sea coast of Thailand. Phuket Marine biological Center Research Bulletin. 59, 43–52.

Christianen, M.J.A., Belzen, J.V., Herman, P.M.J., van Katwijk, M.M., Lamers, L.P.M., van Leent, P.J.M., Bouma, P.M.J., 2017. Low-canopy seagrass beds still provide important coastal protection services. PLoS ONE. 8, e62413

Constanza, R., de Groot, R., Sutton, P., van der Ploeg, S., et al., 2014. Changes in the global value of ecosystem services. Global Environment Change-Human Policy Dimensions. 26, 152–158.

Cunha, A.H., Duarte, C.M., 2005. Population age structure and rhizome growth of *Cymodocea nodosa* in the Ria Formosa (southern Portugal). Mar. Bio. 146: 841–847.

Cullen-Unsworth, L.C., Unsworth, R., 2018. A call for seagrass protection. Science. 361, 6401

Das, H.S., 1996. Status of seagrass habitats of Andaman and Nicobar coast, SACON, Coimbatore, India. Technical Report No. 4

Dorenbosch, M., Grol, MCG, Christianen, M.J. A., Nagelkerken, I., van der Velde, G., 2005. Indo-Pacific seagrass beds and mangroves contribute to fish density and diversity on adjacent coral reefs. Marine Ecology Progress Series. 302, 63–76

Dorenbosch, M., Verberk, W.C.E.P., Nagelkerken, I., van der Velde, G., 2007. Influence of habitat connectivity between fish assemblages of Caribbean seagrass beds, mangroves and coral reefs. Marine Ecology Progress Series. 334, 103–116.

Duarte, C.M., Marbà, N., Agawin, N., Cebrián, J., Enríquez, S. et al., 1994. Reconstruction of seagrass dynamics: age determinations and associated tools for the seagrass ecologist. Mar. Eco. Prog. Ser. 107: 195–209.

Duarte, C.M., Terrados, J., Agawin, N.S.R., Fortes, M.D., Bach, S., Kenworthy, W.J., 1997. Response of a mixed Philippine seagrass meadow to experimental burial. Marine Ecology Progress Series. 147, 285–294

Duarte, C.M., Kennedy, H., Marba, N., Hendriks, I., 2013a. Assessing the capacity of seagrass meadows for carbon burial: current limitations and future strategies. Oceans and Coastal Management. 82, 32–38

Fourqurean, W.J., Duarte, C.M., Marba. N., 2003. Elucidating seagrass population dynamics: Theory, constraints and practice. Limnol. Oceano. 48: 2070–2074.

Gartside. P.S., & Funge-Smith, S., 2013. A review of mangrove and seagrass ecosystems and their linkage to fisheries and fisheries management. Food and Agriculture Organization of the United Nations Regional office for Asia and the Pacific. Bangkok, Thailand, RAP Publication, 74pp.

Guannel, G., Arkema, K., Ruggiero, P., Verutes, G., 2016. The Power of three: Coral reefs, seagrasses and Mangroves protect coastal regions and increase their resilience. PLOS one. 10, 1–22.

Hemminga, M.A., Slim, F.J., Kazungu, J., Ganssen, G.M., Nieuwenhuize, J., Kruyt, N.M., 1994. Carbon outwelling from a mangrove forest with adjacent seagrass beds and coral reefs (Gazi Bay, Kenya). Marine Ecology Progress Series. 106, 209–301.

Howard, J., Hoyt, S., Isensee, K., Telszewski, M., Pidgeon, E., (eds.) (2014). Coastal blue Carbon: Methods for assessing carbon stocks and emissions factors in mangroves, tidal salt marshes, and seagrasses. Conservation International, Intergovernmental Oceanographic Commission of UNESCO, International Union for Conservation of Nature. Arlington, Virginia, USA

Jagtap, T.G., 1991. Distribution of seagrasses along the Indian coast. Aquatic Botany. 40, 379–386

Jagtap, T.G., Komarpant, D.S., and Rodrigues, R.S., 2003. Status of a seagrass ecosystem: an ecologically sensitive wetland habitat from India. Wetlands. 23, 161–170.

Kathiresan, K., and Alikunhi, N.M., 2011. Tropical coastal ecosystems: Rarely Explored for their interaction! Ecologia. 1, 1–22

Kristensen, E., S. Bouillon, T. Dittmar & C. Marchand. 2008. Organic carbon dynamics in mangrove ecosystems: a review. Aquatic Botany 89: 201–219.

Kuo, Y-M., and Lin, H-J., 2010. Dynamic factor analysis of long-term growth trends of the intertidal seagrass Thalassia hemprichii in southern Taiwan. Estuarine, Coastal and Shelf Science. 88, 225–236

Lewis, M.A., and Devereux, R., 2009. Non-nutrient anthropogenic chemicals in seagrass ecosystems: fate and effects. Environment Toxicology and Chemistry. 28:644–661

Lirman D, Cropper WP., 2003. The influence of salinity on seagrass growth, survivorship, and distribution within Biscayne Bay, Florida: field, experimental, and modeling studies. Estuaries. 26: 131–141. doi: 10.1007/BF02691700

McMahon, K.M., Evans, R.D., van Dijk, K., Hernawan, U., Kendrick, G.A., Lavery, P.S., Lowe, R., Puotinen, M., Waycott, M., 2017. Disturbance is an important driver of clonal richness in Tropical seagrasses. Frontiers in Plant Science. 8, 2026.

Medina-Gomez, I., Madden, C.J., Herrera-Silveira, J., Kjerfve, B., 2016. Response of *Thalassia testudinum* morphometry and distribution to Environmental drivers in a Pristine Tropical Lagoon. PLOS one. 1–24. DOI: 10.1371/journal.pone.0164014

Mendoza, A.R.R., Patalinghung, J.M.R., Divinagracia, J.Y., 2019. The benefit of one cannot replace the other: seagrass and mangrove ecosystems at Santa Fe, Bantayan Island. Journal of Ecology and Environment. 43, 1–8

Mishra, A.K., and Mohanraju, R., 2018. Epiphytic bacterial communities in seagrass meadows of oligotrophic waters of Andaman Sea. 5, e4388

Mishra, A.K., and Kumar, M., 2019. Mangrove sediments act as source of nutrients and sink of heavy metals of coastal Andaman Sea. Indian Journal of Geo Marine Sciences. (Accepted article).

Mishra, A.K., Narayana, S., Apte, D., 2019. Physical damage by boat anchors cause loss of Dugong grass (*Halophila ovalis*) population structure. Aquatic Botany (Accepted article)

Nagelkerken, I., M. Dorenbosch, W.C.E.P. Verberk, E. Cocheret de la Morinière & G. van der Velde. 2000. Importance of shallow water biopes of a Caribbean bay for juvenile coral reef fishes: patterns in biope association, community structure and spatial distribution. Marine Ecology Progress Series 202: 175–192.

Nagelkerken, I., Blaber, S.J.M., Bouillon, S., Green, P., Haywood, M., Kirton, L.G., Meynecke, L-O., Pawlik, J., Penrose, H.M., Sasekumar, P., & Somerfield, P.J., 2008. The habitat function of mangroves for terrestrial and marine fauna: a review. Aquatic Botany 89: 155–185

Nobi, E.P., Dilipan, E., Thangaradjou, T., Sivakumar, K., Kannan, L., 2010. Geochemical and geo-statistical assessment of heavy metal concentration in the sediments of different coastal ecosystems of Andaman Islands, India. Estuarine, Coastal and Shelf Science. 87, 253–264

Nobi, E.P., Dilipan, E., Sivakumar, K., and Thangaradjou, T., 2011. Distribution and biology of seagrass resources of Lakshadweep group of Islands, India. Indian Journal of Geo-Marine Science. 40: 624–634.

Nordlund, L.M., Koch, E.W., Barbier, E.B., Creed, J.C., 2016. Seagrass ecosystem services and their variability across genera and geographical regions. Plos One. 10, e0163091.

Nordlund, L.M., Unsworth, R.K., Gullström, M., Cullen-Unsworth, L.C., 2017. Global significance of seagrass fishery activity. Fish Fish: 19, 1–14

Pendleton, L., Donata, M.C., Murray, B.C., Crooks, S., Jenkins, W.A., Sifleet, S., Craft, C., Fourqurean, J.W., Kauffman, J.B., Marbá, N., Megonigal, P., Pidgeon, E., Herr, D., Gordon, D., Baldera, A., 2012. Estimating global “Blue Carbon” emissions from conversion and degradation of vegetated coastal ecosystems. PLOS ONE. 7 (9):e43542

Poovachiranon, S., Chansang, H., 1994. Community structure and biomass of seagrass beds in the Andaman Sea mangrove-associated seagrass beds. Phuket Marine Biological Center Research Bulletin, Thailand. 59, 53–64

Potouroglou, M., Bull, J.C., Krauss, K.W., Kennedy, H.A., Fusi, M., Daffonchio, D., Mangora, M.M., Githaiga, M.N., Diele, K., Huxham, M., 2017. Measuring the role of seagrasses in regulating sediment surface elevation. Scientific Reports. 7, 1–11

Ragavan, P., Saxena, A., Mohan, P.M., Coomar, T., Ragavan, A., 2013. Leaf reddening in seagrasses of Andaman and Nicobar Islands. Tropical Ecology. 54, 269–273.

Ragavan, P., Jayaraj, R.S.C., Muruganantham, M., Jeeva, C., Ubare, V.V., Saxena, A., Mohan, P.M., 2016. Species composition and Distribution of Seagrasses of the Andaman and Nicobar Islands. VEGETOS. 29, 78–87.

Rattanachot, E., Stankovis, M., Aongsara, S., Prathep, A., 2018. Ten Years of conservation efforts enhance seagrass cover and carbon storage in Thailand. Aquatic Botany. 61, 441–451

Rietkerk, M., S. C. Dekker, P. C. de Ruiter & J. van de Koppel, 2004. Self-organized patchiness and catastrophic shifts in ecosystems. Science 305: 1926–1929

Rollon, R., Cayabyab, N.M., Fortes, M., 2001. Vegetative dynamics and sexual reproduction of monospecific Thalassia hemprichii meadows in the Kalayaan Island group. Aquatic Botany. 71, 239–246

Saunders, M.I., Atkinson, S., Klein, C.J., Weber, T., Possingham, H.P., 2017. Increased sediment loads can cuase non-linear decrease in seagrass suitable habitat extent. 12, e0187284

Saxena, A., Ragavan, P and Ambika, R., 2010. Assessment of status and diversity of seagrasses in Andaman and Nicobar Islands. In Proceedings of the National Seminar on Tropical Ecosystems: Structure, Function and Services, IFGTB, Coimbatore, pp. 31–38

Saenger, P., Gartside, D., Funge-Smith, S., 2012. A review of mangrove and seagrass ecosystems and their linkage to fisheries and fisheries management. FAO Regional Office for Asia and the Pacific, Bangkok, Thailand, RAP Publication, 2013/09, 74pp.

Sahu, S.C., Suresh, H.S., Murthy, I.K., Ravindranath, N.H., 2015. Mangrove Area assessment in India: Implications of loss of mangroves. Journal of Earth Science and Climate Change. 6, 280.

Savurirajan, M., Lakra, R.K., Equbal, J., Ganesh, K.S.T., 2018. A note on morphometric, shoot density and biomass of Thlassia hemprichii from the south Andaman coast, Andaman and Nicobar Islands, India. Indian Journal of Geo Marine Sciences. 47, 1222–1227

Short, F.T., Coles, R.C., 2001. Global Seagrass Research Methods. Elsevier Science, B.V., Amsterdam, 473 pp.

Short, F.T., and Duarte, C.M., 2001. Methods for the measurement of seagrass growth and production. In eds., Global Seagrass Research Methods. Elsevier Science B.V., Amsterdam. 473pp.

Short, F.T., Carruthers, T., Dennison, W., Waycott, M., 2007. Global seagrass distribution and diversity: A bioregion model. Journal of Experimental Marine Biology and Ecology. 350, 3–20.

Short, F.T., Polidoro, B., Livingstone, S.R., Carpenter, K.E., et al., 2011. Extinction risk assessment of the world’s seagrass species. Biological Conservation. 7, 1961–1971.

Smith, N.M., and Walker, D.I., 2002. Canopy structure and pollination biology of the seagrasses *Posidonia australis* and *P. sinuosa* (Posidoniaceae). Aquatic Botany. 74, 57–70

Sokal, R. R. and Rohlf, F.J., 2012. Biometry: the principles and practice of statistics in biological research. 4th edition. W. H. Freeman and Co.: New York. 937 pp.

Spalding, M.D., Taylor, M.L., Ravilious. C, Short, F., Green. E. Global overview: The distribution and status of seagrasses. In: Green E, Short FT, editors. World atlas of seagrasses. Berkeley: University of California Press; 2003. pp. 5–26

Thangaradjou, T., Nobi, E.P., Dilipan, E., Sivakumar, K., Susila, S. 2010. Heavy metal enrichment in seagrasses of Andaman Islands and its implication to the health of the coastal ecosystem. Indian Journal of Geo Marine Science. 39, 85–91.

Thangaradjou, T., Nobi, E.P., Dilipan, E., 2010. Distribution of seagrasses along the Andaman and Nicobar Islands: A post Tsunami survey. Recent Trends in Biodiversity of Andaman and Nicobar Islands, 157–160.

Thangaradjou, T., Nobi, E.P., Dilipan, E., Sivakumar, K., Kannan, L., 2009. Threats to seagrass under of India, Seagrass Watch. 39, 20–21

Tussenbroek, B.I., Vonk, J.A., Stapel, J., Erftemeijer, P.L.A., Middleburg, J.J., Zieman, J.C., 2006. The biology of Thalassia: Paradigms and Recent Advances in Research. In. Seagrass: Biology, ecology and Conservation. Pp-409–439. Springer

Wang, M., Z. Huang, F. Shi & W. Wang. 2009. Are vegetated areas of mangroves attractive to juvenile and small fish? The case of Dongzhaigang Bay, Hainan Island, China. Estuarine, Coastal and Shelf Science 85: 208–216.

